# Taste quality and hunger interactions in a feeding sensorimotor circuit

**DOI:** 10.1101/2022.03.06.483180

**Authors:** Philip K. Shiu, Gabriella R. Sterne, Stefanie Engert, Barry J. Dickson, Kristin Scott

## Abstract

Taste detection and hunger state dynamically regulate the decision to initiate feeding. To study how context-appropriate feeding decisions are generated, we combined synaptic resolution circuit reconstruction with targeted genetic access to specific neurons to elucidate a gustatory sensorimotor circuit for feeding initiation in adult *Drosophila melanogaster*. This circuit connects gustatory sensory neurons to proboscis motor neurons through three intermediate layers. Most neurons in this pathway are necessary and sufficient for proboscis extension, a feeding initiation behavior, and respond selectively to sugar taste detection. Pathway activity is amplified by hunger signals that act at select second-order neurons to promote feeding initiation in food-deprived animals. In contrast, the feeding initiation circuit is inhibited by a bitter taste pathway that impinges on premotor neurons, illuminating a local motif that weighs sugar and bitter taste detection to adjust behavioral outcome. Together, these studies reveal central mechanisms for the integration of external taste detection and internal nutritive state to flexibly execute a critical feeding decision.

## Introduction

The decision to initiate feeding depends both on the quality of available food and current nutrient needs. The gustatory system detects nutritious and noxious compounds in the environment and evaluates food quality. Food quality information is integrated with internal nutritive state to ensure that food intake matches energy demands. How do central neural circuits evaluate taste information in the context of internal nutritive state to make feeding decisions? As feeding decisions are universal and essential for survival, animals as diverse as humans and *Drosophila* share similar strategies to detect taste compounds in the environment and assess nutrient needs. Peripheral taste detection in mammals and insects is mediated by sensory cells that detect specific taste modalities and elicit innate feeding behaviors. Both mammals and flies have sugar-, bitter-, water-, and salt-sensing gustatory cells (Liman, et al., 2014; Yarmolinsky et al., 2009). Activation of sugar-sensing gustatory cells triggers feeding initiation, whereas activation of bitter-sensing cells inhibits feeding. Mammals and insects also evaluate internal nutrient needs with similar strategies (Augustine et al., 2018; Leopold and Perrimon, 2007; Pool and Scott, 2015; Nässel and Zandawala, 2019). Neuromodulators released from neurosecretory centers and the gut signal hunger or satiety to oppositely regulate feeding. Disruption of these hunger and satiety signals results in obesity and anorexia in mammals and insects.

Although gustatory sensory neurons have been shown to be modulated by hunger signals and conflicting taste information (Chu et al., 2014; French et al., 2015; Inagaki et al., 2012; Inagaki et al., 2014; Jeong et al., 2013; Meunier et al., 2003), central mechanisms that modulate feeding decisions are unclear because the identity, structure, and function of central feeding initiation circuits is unknown. Recent advances in brain-wide synaptic connectivity mapping (Eckstein et al., 2020; Dorkenwald et al., 2020; Zheng et al., 2018) and precise genetic access to single neurons (Luan et al., 2006; Dionne et al., 2018) make *Drosophila melanogaster* an ideal system to interrogate how the central brain computes feeding decisions. Taste detection in adult *Drosophila* begins with activation of gustatory receptor neurons (GRNs) found in sensory structures located on the body surface, including the external mouthparts, or proboscis labellum (Dethier, 1976; Stocker, 1994; Montell, 2021; Scott, 2018). The axons of proboscis GRNs project to the primary taste and premotor center of the insect brain, the subesophageal zone (SEZ) (Stocker and Schorderet, 1981; Kendroud et al., 2018; Miyazaki and Ito, 2010). The motor neurons that execute feeding have cell bodies and dendrites in the SEZ near GRN axons, suggesting a local feeding circuit (Gordon and Scott, 2009; McKellar et al., 2020; Schwarz et al., 2018). However, only a few isolated interneurons have been implicated in feeding initiation (Flood et al., 2013; Kain and Dahanukar, 2015).

To investigate how neural circuits transform taste detection into context-appropriate feeding decisions, we combined electron microscopy-based circuit reconstruction, genetic tools that provide access to single cell types, optogenetics, and imaging of taste responses in awake, behaving animals to uncover a circuit-level view of feeding initiation in *Drosophila melanogaste*r. This work delineates the neural circuit that transforms taste detection into the motor actions of feeding initiation from sensory inputs to motor outputs and reveals central mechanisms that integrate taste detection with internal physiological state to shape behavior.

## Results

### Gustatory receptor neurons synapse onto multiple second-order neurons

To examine neural circuits for feeding initiation, we identified neurons directly postsynaptic to gustatory sensory axons in the central brain. We utilized the full adult fly brain (FAFB) electron microscopy (EM) volume (Zheng et al., 2018) to manually reconstruct neurons postsynaptic to 17 labellar GRN axons that likely correspond to sugar-sensing GRNs (Engert et al., 2021). Fifteen second-order taste neurons and their synapses were fully reconstructed (Figures 1A, 1B and S1) in the CATMAID platform (Saalfeld et al., 2009). The manually reconstructed neurons were compared with those in the recently released Flywire dataset, a dense, machine learning based reconstruction of FAFB neurons (Eckstein et al., 2020; Dorkenwald et al., 2020), revealing that they represent 12 of the 13 cell types with the most synapses from candidate sugar GRNs (Figure S1F). These second-order neurons have not previously been characterized, except for G2N-1, which was identified as a candidate second-order gustatory neuron based on anatomical proximity to sugar-sensing GRNs (Miyazaki et al., 2015). Each of the second-order neurons is a local SEZ interneuron with arbors that overlap extensively with sugar GRN termini.

**Figure 1.**
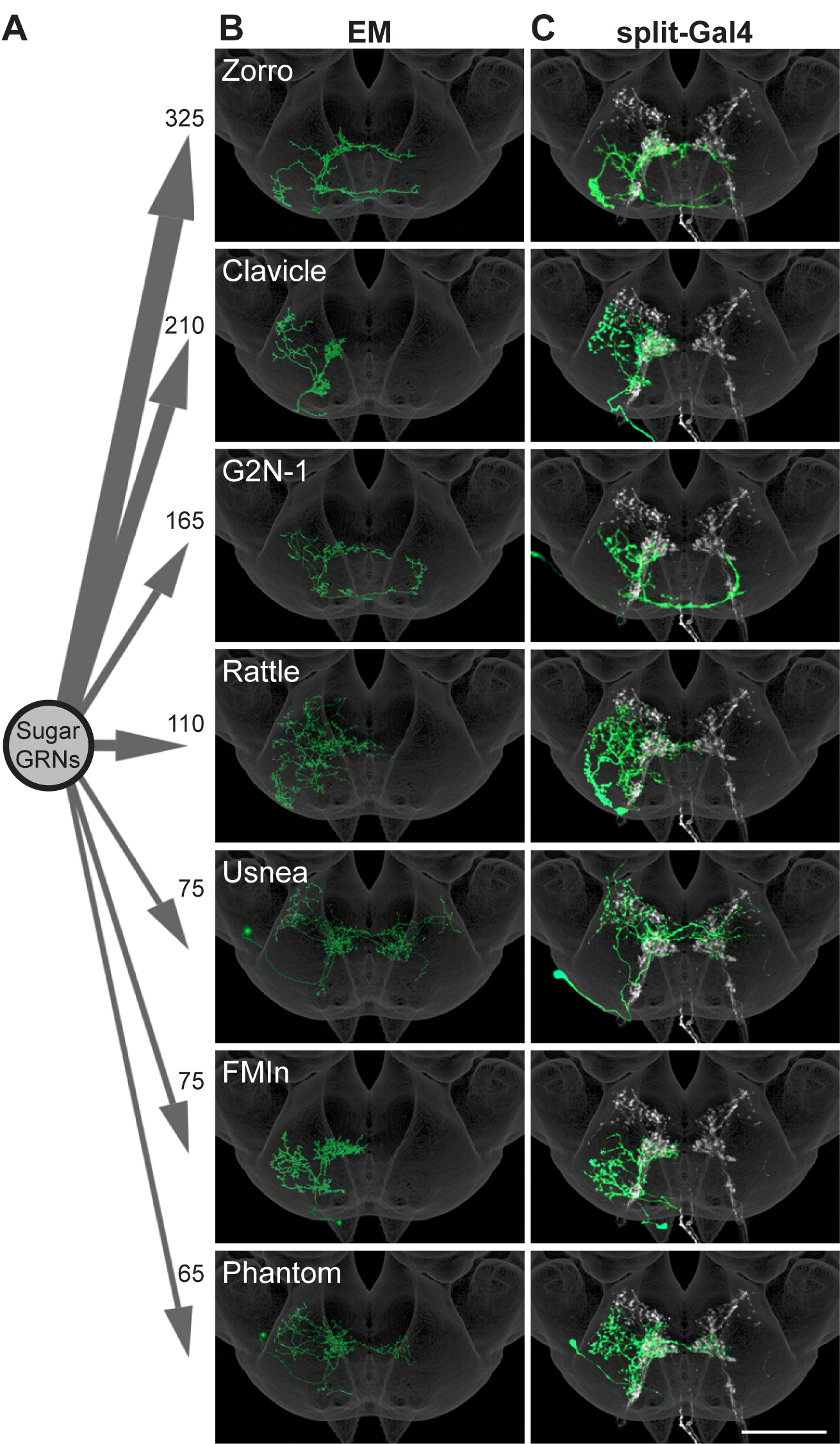
Sugar-sensing GRNs synapse onto multiple second-order neurons. (A) Aggregate synaptic connectivity from sugar GRNs onto second-order sugar neurons. Numbers indicate the total number of synapses that the 17 candidate sugar GRNs make onto each second-order neuron. (B-C) Manually reconstructed EM skeletons (B) and registered neural images in split-Gal4 lines (C) for each second-order neuron in the subesophageal zone of the *Drosophila* brain. Sugar GRNs are depicted in white, JRC 2018 unisex coordinate space shown in gray (C). Scale bar is 50 μm. See also Figure S1 for EM reconstructions of additional second-order neurons and synaptic connectivity counts.

### Multiple second-order taste neurons influence proboscis extension

To test whether second-order gustatory neurons participate in feeding behaviors, we identified split-Gal4 lines that provide specific genetic access to individual second-order cell types, using NBLAST comparisons (Costa et al., 2016) to a library of SEZ split-Gal4 lines (Sterne et al., 2021). This provided split-Gal4 matches for 7 second-order neurons (Figure 1C and S2A). Additionally, we used intersectional approaches to gain genetic access to two additional second-order neurons, Cleaver (Figure S2A) and Zorro (Figure 1C). The split-Gal4 lines are exquisitely specific for each of the 9 second-order gustatory neurons, providing the opportunity to evaluate their function.

As activation of sugar-sensing GRNs on the proboscis labellum causes the fly to extend its proboscis to initiate feeding (Dethier, 1976), we tested whether activation or inhibition of second-order taste neurons influences this behavior. We expressed the red-shifted channelrhodopsin CsChrimson (Klapoetke et al., 2014) selectively in each second-order taste neuron, activated each with 635 nm light, and examined the proboscis extension response (PER). Remarkably, optogenetic activation of 7 of the 9 second-order taste neurons elicited proboscis extension (Figures 2A and S2B).

**Figure 2.**
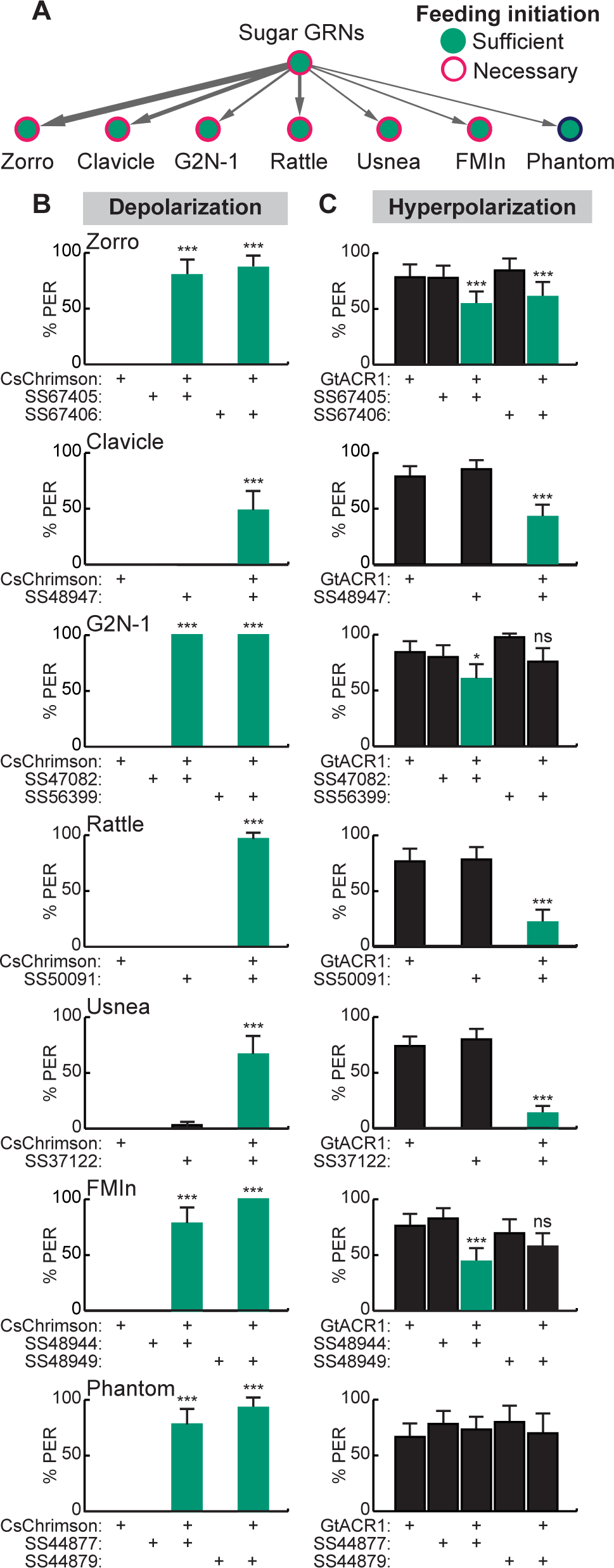
Activation and inhibition of second-order taste neurons influences proboscis extension. (A) Schematic of proboscis extension phenotypes of second-order neurons. Line thickness corresponds with total synaptic connectivity. (B) CsChrimson-mediated activation of seven second-order neurons elicits proboscis extension. n=30 flies per genotype. (C) GtACR1-mediated inhibition of second-order neurons reduces proboscis extension to 50 mM sucrose. n= 46-83 flies per genotype. (B-C) The fraction of flies exhibiting PER upon optogenetic or 50 mM sucrose stimulation. Mean ± 95% confidence interval (CI), Fisher’s Exact Tests, *p<0.05, ***p<0.001. See also Figure S2 for additional PER phenotypes of second-order sugar neurons.

Moreover, inhibiting the activity of each second-order neuron individually, by optogenetic activation of the anion channelrhodopsin GtACR1 (Mohammad et al., 2017) reduced proboscis extension to 50 mM sucrose in food-deprived flies, for six of the seven second-order neurons that elicited PER upon activation (Figure 2B). At a higher sucrose concentration (100 mM), neural inhibition of only two of the second-order neuron classes decreased proboscis extension (Figure S2C). These studies argue that multiple second-order neurons contribute to normal feeding initiation behavior and suggest that the partial redundancy of these second-order neurons ensures robust feeding.

### Second-order taste neurons activate a local SEZ circuit for feeding initiation

How does activation of a diverse set of second-order neurons drive proboscis extension? Previous studies have demonstrated that proboscis motor neuron 9 (MN9) contracts the rostrum protractor muscle to extend the proboscis and is required for feeding initiation (Gordon and Scott; McKellar et al., 2020). We located MN9 in the FAFB EM volume, by examining large SEZ neurons that lack synaptic output. To identify a pathway from taste detection to proboscis extension, we reconstructed presynaptic partners of MN9 and postsynaptic partners of second-order taste neurons.

This strategy identified a minimal pathway from taste detection to proboscis extension, composed of interconnected second-order neurons, third-order neurons each receiving inputs from a subset of second-order neurons, and feedforward premotor neurons (Figure 3A and S3A). The third-order neurons represent a small subset based on comparisons to Flywire automated reconstructions (Figure S3B). They include one previously characterized neuron, the putative feeding command neuron, Fdg (Flood et al., 2013) (Figure 3B) and a set of descending neurons, Bract, that project to the ventral nerve cord (Sterne et al., 2021). The premotor neurons are strongly connected to MN9, representing approximately 13% of the synaptic input onto MN9 (Figure S3C). There are direct connections between three second-order neurons (G2N-1, Zorro and FMIn) and pre-motor neurons, and additional paths via third-order neurons to premotor neurons.

**Figure 3.**
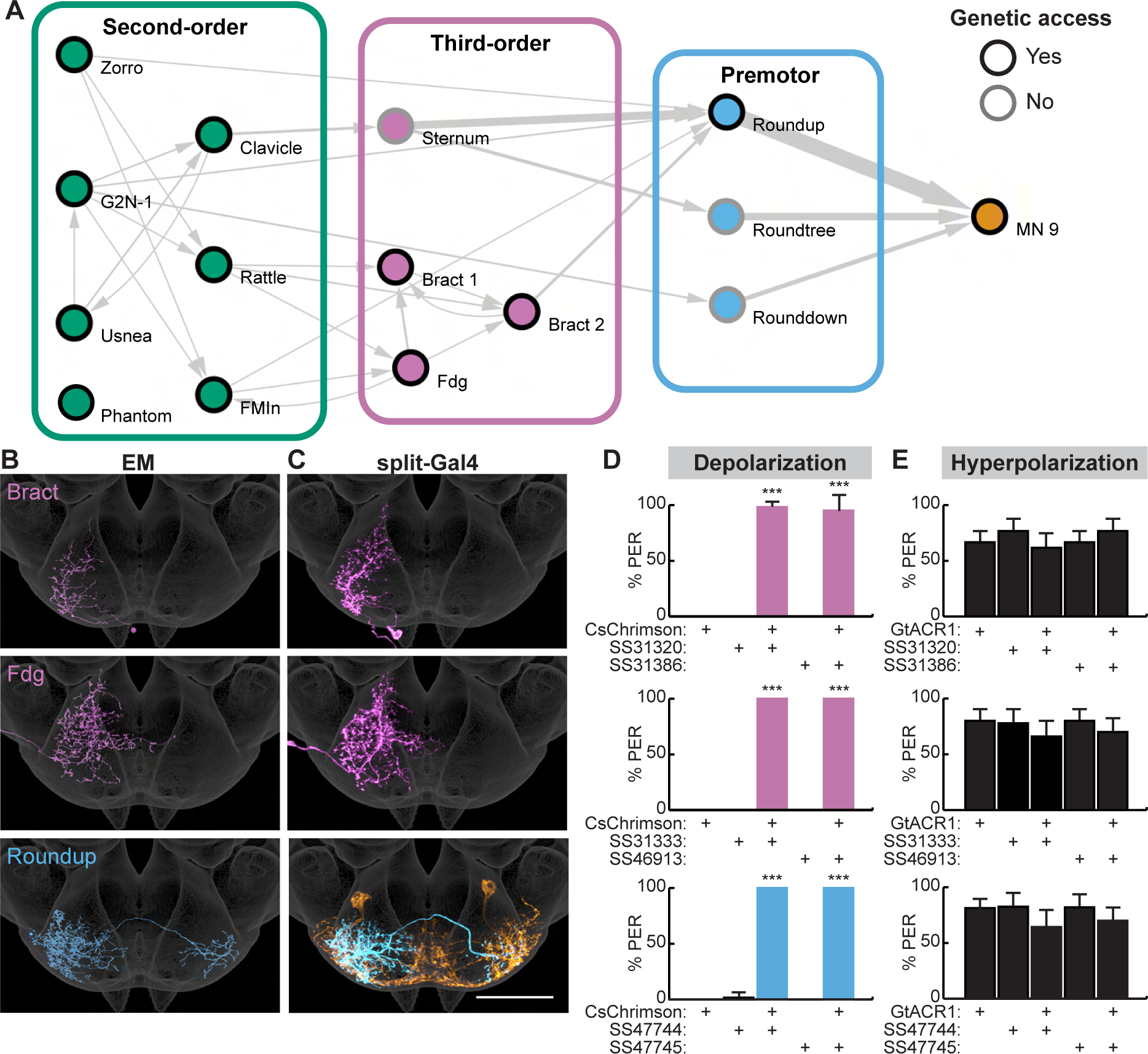
Second-order neurons synapse onto a local sensorimotor circuit for feeding initiation. (A) Schematic of the feeding initiation circuit. Circles outlined in black denote neurons with split-Gal4 genetic access, circles with gray outlines denote neurons without split-Gal4 genetic access. Line thickness represents synaptic connectivity of more than 5 synapses. (B-C) EM neural reconstructions (B) and registered neural images in split-Gal4 lines (C) of third-order or premotor neurons in the SEZ. Scale bar is 50 μm. JRC 2018 unisex coordinate space shown in gray, MN9 morphology shown in orange. (D) Cs-Chrimson-mediated activation of third-order or premotor neurons elicits PER. n=30 flies per genotype. (E) GtACR1-mediated inhibition of third-order or premotor neurons does not influence PER to 50 mM sucrose. n=40-70 flies per genotype. (D-E) Mean ± 95% CI, Fisher’s Exact Tests, ***p<0.001. See Figure S3 for synaptic counts.

To investigate the function of deeper layers of this circuit, we identified split-Gal4 lines that selectively label two third-order neurons and one premotor neuron (Figure 3C) using NBLAST comparisons with SEZ split-Gal4 lines (Sterne et al., 2021). Optogenetic activation of third-order or premotor neurons with CsChrimson revealed that each cell type elicits robust proboscis extension (Figure 3D). However, acute inhibition of the third-order or premotor neurons with GtACR1 did not influence PER to 50 mM sucrose (Figure 3E), consistent with multiple pathways to proboscis motor neurons. Thus, by combining EM tracing studies with precise neural manipulations afforded by split-Gal4 lines, we have elucidated a neural circuit that promotes feeding initiation upon sweet taste detection.

### Feeding initiation neurons respond to sugar taste detection

To examine how taste information is processed by the feeding initiation circuit to guide feeding decisions, we monitored taste-induced activity of each neuron in the circuit. The proboscis was stimulated with water, sugar, or bitter taste solutions, while monitoring GCaMP6s calcium activity (Chen et al., 2013) in live flies (Harris et al., 2015). Eight of the ten neural classes responded to sugar taste presentation in food-deprived animals, and not to water or bitter solutions (Figure 4). Two second-order cell types responded to water taste detection: Usnea responded specifically to water and Phantom responded equally to water and sugar detection. Usnea and Phantom are reciprocally connected with GRNs (Figure S1D), suggesting that these second-order cell types may tune GRN responses in the presence of water. One third-order neuron, Fdg, did not respond to proboscis taste stimulation, but did respond to optogenetic activation of sugar-sensing GRNs (Figure S4A), suggesting that it may respond to pharyngeal or leg taste detection. Together, these studies reveal that sugar taste is processed by a multi-layered neural circuit to initiate feeding in hungry flies.

**Figure 4.**
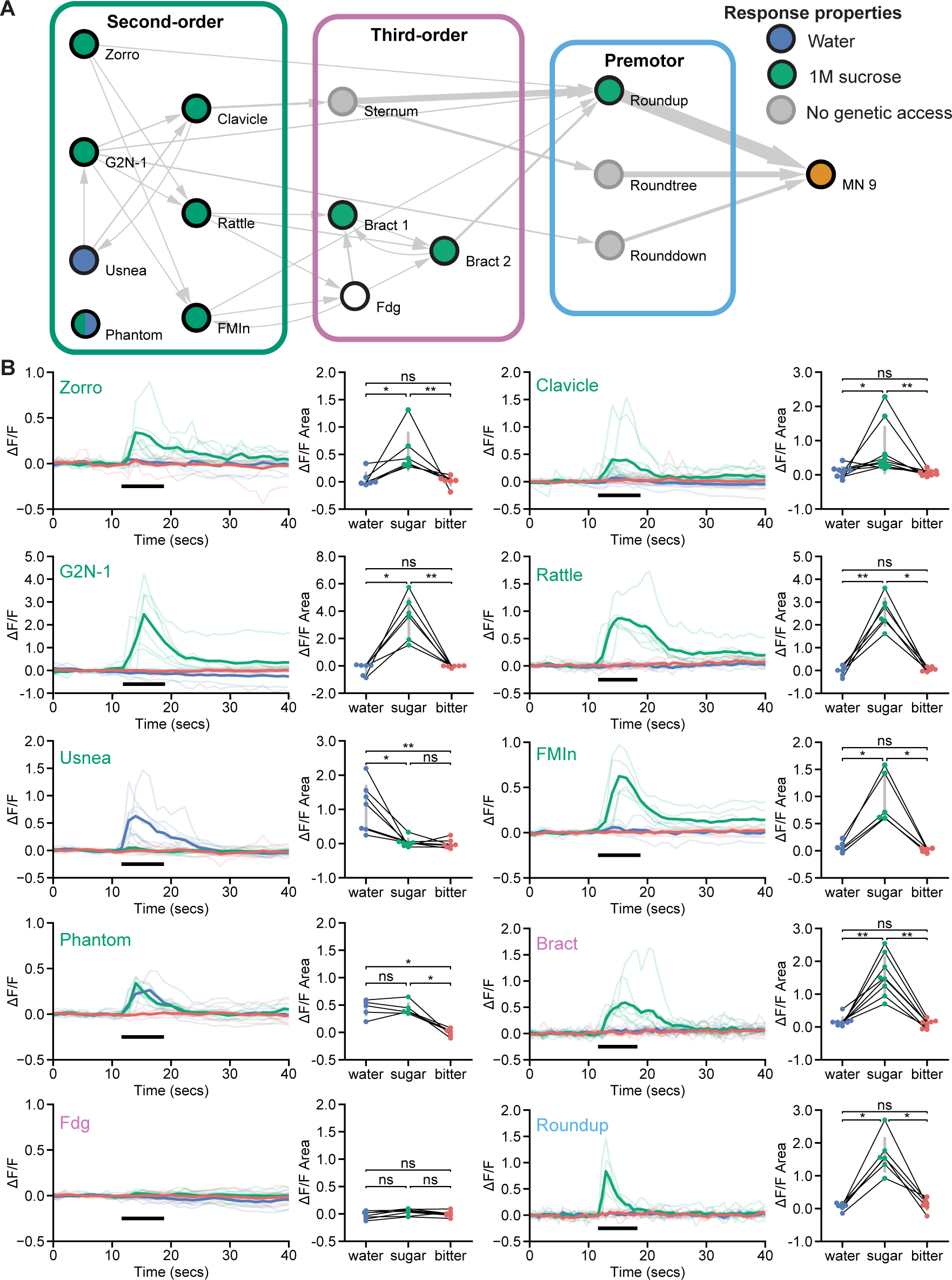
Feeding initiation neurons respond to taste detection. (A) Connectivity schematic of the feeding initiation circuit, where filled green circles represent cell types that respond to sugar detection, while filled blue circles represent cell types that respond to water detection. One cell type, Phantom, responds to both sugar and water (split blue and green circle). Fdg did not respond to proboscis taste detection (white circle), but see Figure S4A for responses to optogenetic activation of sugar GRNs. (B) Calcium responses of feeding initiation neurons to stimulation of the proboscis in food- deprived flies. For each cell type, GCaMP6s fluorescence traces are shown on the left of the panel (ΔF/F), while ΔF/F area for each trace is shown on the right, with thin black lines indicating sample pairing. The proboscis of each tested individual was stimulated with water (green), sugar (blue), and bitter (red) tastants in sequential trials during the indicated period (thick black line). The following split-GAL4 lines were imaged for each cell type: Clavicle; SS48947, FMIn; SS48944, Zorro; SS67405, G2N-1; SS47082, Usnea; SS37122, Phantom; SS68204, Rattle; SS50091, Fdg; SS31345, Bract; SS31386, Roundup; SS47744. Quade’s Test with Quade’s All Pairs Test, using Holm’s correction to adjust for multiple comparisons, ns p>0.05, *p≤0.05, **p≤0.01. See also Figure S4 for addtional calcium imaging studies of feeding initiation neurons.

To test whether responses in the proboscis extension circuit are altered based on specific nutrient need, we examined taste responses in flies that were thirsty rather than hungry. High hemolymph osmolality is a key signal of thirst that acts on central neurons to promote water consumption (Jourjine et al., 2016). We mimicked a thirsty-like state by increasing hemolymph osmolality, which enhanced water responses in water-sensing GRNs (Figure S3B). In four of the five central neurons tested, response profiles were similar in food-deprived and thirsty-like flies (Figure 4 and S4C). However, one second-order neuron, Clavicle, responded to water and to sugar taste detection in a thirsty-like state but only to sugar in a hungry state (Figure 4 and S4C). These results suggest that state-dependent responses to water at a single node (Clavicle) may tune the responsiveness of the pathway to bias acceptance of more dilute sugar solutions.

However, the majority of neurons uncovered here are dedicated to sugar taste detection regardless of whether the animal is hungry or thirsty.

### Feeding initiation is modulated by hunger at specific nodes

How is sugar taste information integrated with hunger state to promote feeding initiation in food-deprived flies? Hunger modulates sugar GRN activity (Inagaki et al., 2012); however, whether sensory gating is the only mechanism for hunger regulation or whether modulation of central neurons contributes to an altered network state in hungry animals has not been examined. To comprehensively investigate how taste detection is integrated with hunger state to initiate feeding, we optogenetically activated each neuron in the PER circuit in either fed or food-deprived flies and examined behavior.

Optogenetic activation has the advantage of bypassing changes in sugar sensory detection that propagate through the circuit, enabling the evaluation of central circuit changes.

We reasoned that activating neurons upstream or at the node(s) where hunger modulation occurs will cause differences in proboscis extension rates between hungry and fed flies, whereas activating neurons beyond the site where hunger impinges will not. Indeed, CsChrimson-mediated activation of primary sugar-sensing neurons caused higher proboscis extension rates in food-deprived flies compared to fed flies, whereas activation of MN9 elicited the same proboscis extension rate in food-deprived and fed flies (Figure 5A - 5C). Moreover, activation of two second-order neurons, G2N-1 and Clavicle, increased proboscis extension in food-deprived flies, whereas activation of all other neural classes did not (Figure 5A and 5D). Thus, hunger signals act on sensory neurons to increase detection sensitivity and on a specific set of second-order interneurons to amplify sugar pathway activation and promote feeding.

**Figure 5.**
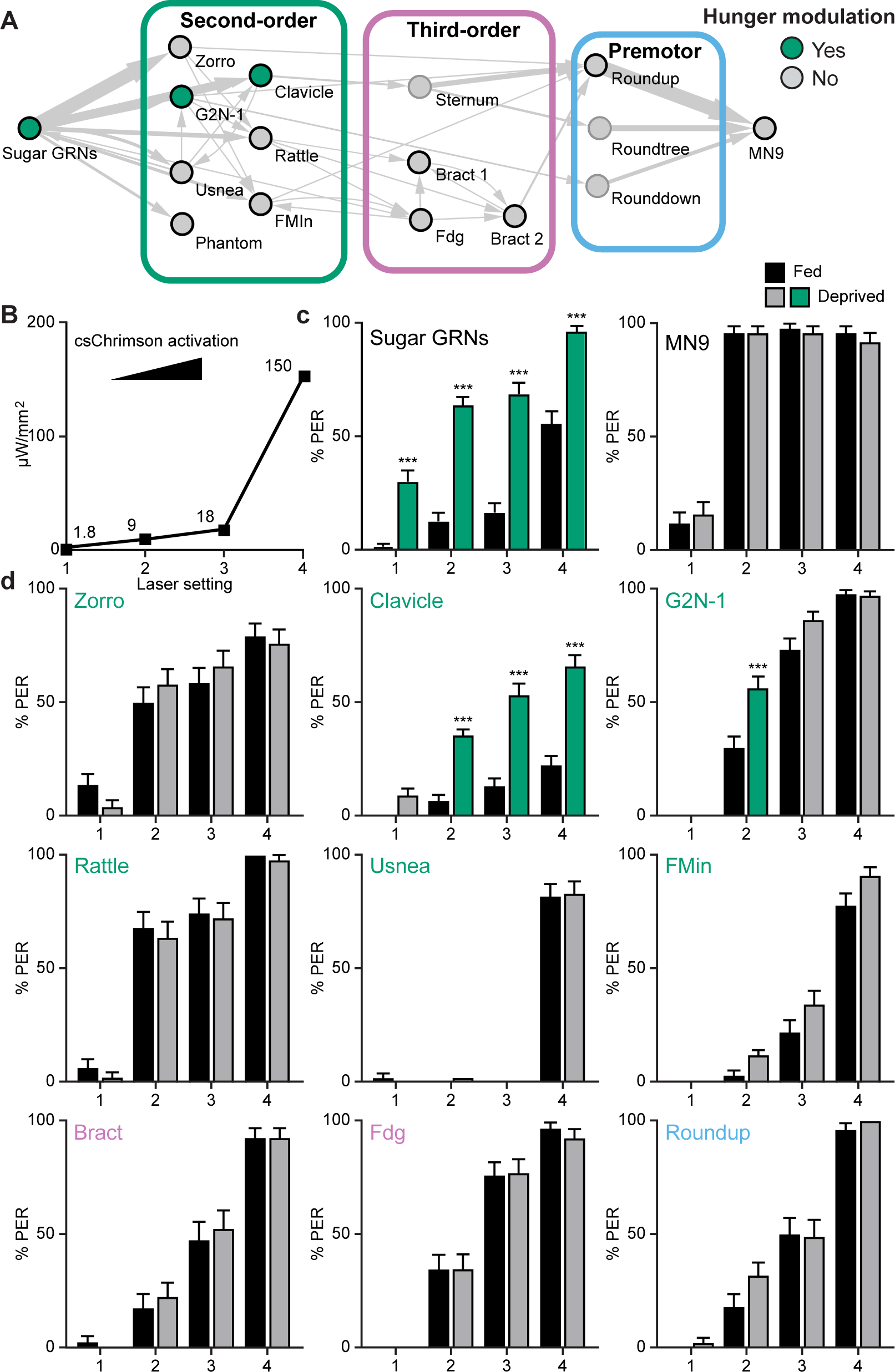
Hunger acts on a subset of second-order central neurons to modulate behavior. (A) Schematic of the feeding initiation circuit, with filled green circles representing nodes that are hunger-modulated. (B) Optogenetic activation at four different light intensities. (C) Activation of sugar-sensing neurons results in different feeding initiation rates between fed and food-deprived flies (left) whereas activation of MN9 does not (right), at four different light intensities. n=50. (D) Optogenetic activation of second-order, third-order, and premotor neurons in either fed or food-deprived flies. n= 39-103. Mean ± 95% CI, Fisher’s Exact Tests, ***p<0.001.

### Premotor neurons integrate sweet and bitter taste information

Animals evaluate both internal nutritive state and food quality to decide whether to initiate feeding. To investigate how food quality alters feeding initiation, we examined how the detection of bitter compounds is integrated with sugar taste information in the feeding initiation circuit. Previous studies have demonstrated that bitter compounds inhibit sugar-sensing gustatory neurons to prevent feeding (Chu et al., 2014; French et al., 2015; Jeong et al., 2013; Meunier et al., 2003), but have not addressed how downstream neural circuitry modulates appetitive feeding behaviors in response to bitter taste detection. To investigate central mechanisms of bitter modulation, we examined whether pathways from bitter GRNs intersect with the feeding initiation pathway.

As bitter GRNs do not directly synapse with neurons in the feeding initiation circuit, we asked whether second-order bitter neurons synapse onto the feeding initiation pathway. We reconstructed neurons downstream of bitter GRNs (Engert et al., 2021) in the EM volume and identified a second-order bitter neuron, Scapula, which receives over 150 synapses from bitter GRNs and is the second-most strongly connected cell type with bitter GRNs (Figure 6A and S5). Scapula synapses directly onto two feeding initiation premotor neurons, Roundup and Rounddown, but not onto second-order appetitive taste neurons.

**Figure 6.**
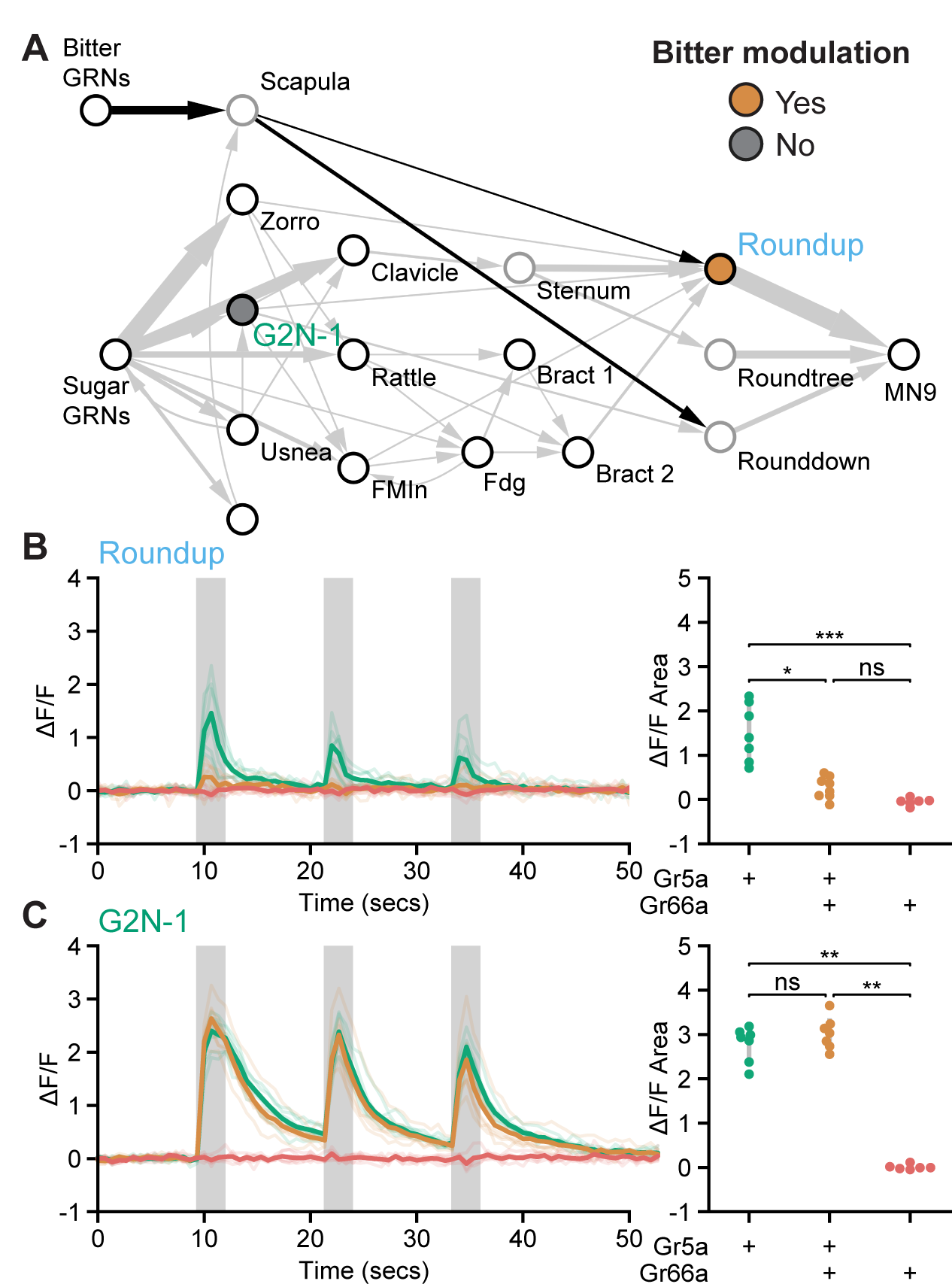
Premotor neurons integrate sweet and bitter taste detection. (A) Schematic of the feeding initiation circuit, showing a pathway from bitter GRNs to premotor neurons. Filled maize circle labels a premotor neuron inhibited by bitter tastants, filled gray circle labels an upstream second-order neuron that is not inhibited by bitter tastants. (B and C) Calcium responses of feeding circuit neurons to optogenetic activation of sugar (green, Gr5a-LexA), sugar plus bitter (maize, Gr5a-LexA plus Gr66a- LexA), or bitter (red, Gr66a-LexA) GRNs in food-deprived flies. For each cell type, Syt::GCaMP7b fluorescence traces are shown on the left of the panel (ΔF/F), while ΔF/F area for each trace is shown on the right. Periods of stimulation with 660 nm light are indicated with vertical gray bars. (B) SS47744 was imaged to examine Roundup responses. (C) SS47082 was imaged to examine G2N-1 responses. Kruskal Wallace test with Dunn’s test using Holm’s correction to adjust for multiple comparisons, ns p>0.05, *p≤0.05, **p≤0.01. See Figure S5 for synaptic counts of second- order bitter neurons.

Because bitter taste detection inhibits proboscis extension, we hypothesized that Roundup and Rounddown would be inhibited by Scapula to prevent proboscis extension to sugar in the presence of bitter compounds. To test this, we monitored activity in Roundup by *in vivo* calcium imaging while activating sugar GRNs, bitter GRNs, or both, using optogenetics to bypass sensory modulation. Roundup responded to optogenetic activation of sugar GRNs but not bitter GRNs (Figure 6B), as expected based on its response to taste compounds (Figure 4B). Upon co-activation of sugar and bitter GRNs, the Roundup response was dramatically decreased compared to the response to sugar GRN activation alone, arguing that bitter signals suppress the feeding initiation pathway. To test whether this bitter suppression reflects a central mechanism acting at Roundup, we monitored activity in a second-order neuron, G2N-1, directly upstream of Roundup, and found that its response upon co-activation of sugar and bitter GRNs was indistinguishable from its response to sugar GRN activation alone (Figure 6C).

Together, the EM and imaging studies demonstrate that sugar and bitter tastes are integrated at feeding initiation premotor neurons, providing a central mechanism to reject sweet foods laced with bitter compounds.

## Discussion

In this study, we coupled EM circuit reconstruction with the ability to precisely monitor and manipulate single neurons to elucidate how a complex nervous system orchestrates the decision to initiate feeding. First, we delineate the sensorimotor circuit for feeding initiation from sensory inputs to motor outputs with cellular and synaptic resolution. Then, we demonstrate how this central circuit integrates taste detection with internal state, providing mechanistic insight into how taste modalities and feeding decisions are encoded in the brain.

### A local, interconnected network transforms sweet taste detection into behavior

Previous studies in *Drosophila* have identified gustatory neurons, motor neurons, and three candidate interneurons that influence feeding initiation (Gordon and Scott, 2009; McKellar et al., 2020; Flood et al., 2013; Kain and Dahanukar, 2015; Miyazaki et al., 2015; Talay et al., 2017). Here, by elucidating a complete sensorimotor circuit with synaptic resolution, we provide a comprehensive view of the neural pathway that elicits proboscis extension, the first step in feeding. The feeding initiation pathway is a local circuit, with three- and four- synaptic relays to motor output. All neurons elicit proboscis extension upon optogenetic activation, showing the tight link of all relays to behavior.

Inhibiting activity of single second-order neurons reduced the behavioral response whereas inhibiting activity of third-order or premotor neurons did not. This partial redundancy is consistent with the circuit architecture, revealing multiple paths from second-order neurons to premotor neurons. This partial redundancy may enable proboscis extension to be recruited in different contexts to ensure robust feeding.

Most neurons in this pathway respond to sugar taste detection, but not water or bitter tastes, in food-deprived flies, demonstrating a direct line from sweet taste detection to the motor output for feeding. How water taste modulates proboscis extension in thirsty flies will require further study. Importantly, we identified and characterized three second-order neurons that respond to water taste detection: one is selective for water taste, another responds to both water and sugar tastes, and a third shows state-dependent sugar and water taste responses. Further study of these second-order neurons and their connectivity will be critical to evaluate the degree of separation or convergence of water and sugar pathways for feeding initiation.

Of the interneurons identified here, only G2N-1 and Fdg have previously been implicated in feeding. G2N-1 was identified as a candidate sugar-sensing second-order neuron based on its anatomical proximity to gustatory axons alone (Miyazaki et al., 2015); here, we elucidated its functional role in taste detection and feeding initiation.

Fdg was isolated as a “feeding command neuron”, able to elicit multiple steps in feeding, including proboscis extension (Flood et al., 2013). In our calcium imaging studies, Fdg did not respond to proboscis taste stimulation but did respond to optogenetic activation of sugar GRNs. This suggests that Fdg may receive gustatory signals from GRNs on the pharynx or legs. Our studies demonstrate that Fdg is a third-order neuron in the feeding initiation pathway, with synaptic connections to Bract descending neurons. A description of the reconstruction of all FlyWire DNs is in preparation (K. Eichler, M. Costa, G. Card, G. Jefferis, personal communication). As Bract synapses with proboscis premotor neurons and ventral cord circuits, Fdg and Bract are well-poised to coordinate proboscis extension with other steps in feeding.

The architecture of the circuit provides a platform to investigate how taste signals are transformed in the brain to drive behavior. In this study, we focused on MN9, the rostrum protractor motor neuron that elicits proboscis extension, as a key readout of proboscis extension behavior. However, proboscis extension involves not only rostrum protraction but also extension of the haustellum and opening of the labellum, controlled by additional motor neurons (McKellar et al., 2020). We hypothesize that the connectivity among second-order and third-order neurons may coordinate the precise temporal activation of different muscle groups for coordinated extension. Moreover, proboscis extension is followed by ingestion then meal termination (Dethier, 1976; Pool and Scott, 2014). Continued expansion and exploration of this pathway will provide the opportunity to examine how different feeding subprograms are timed and coordinated to elicit feeding in natural environments.

### Hunger tunes second-order neurons to promote sugar responses

Studies in *C. elegans*, *Drosophila*, and mammals have demonstrated that a key site of hunger regulation is at the peripheral chemosensory neurons, altering sensitivity of detection (Chalasani et al., 2010; Kawai et al., 2000; Root et al., 2011; Savigner et al., 2009; Sengupta, 2013). For example, dopamine enhances the sensitivity of *Drosophila* sugar-sensing gustatory neurons to promote proboscis extension at lower sucrose concentrations (Inagaki et al., 2012; Marella et al., 2012). Hunger modulation of taste processing beyond sensory neurons has been more challenging to evaluate, both because of lack of knowledge of central networks and because changes at the sensory level propagate through the network.

To isolate the role of central brain neurons in hunger modulation, we used the precise genetic access available in *Drosophila* to activate each node of the feeding initiation pathway and examined the behavioral response elicited in fed and food-deprived flies. These studies pinpoint the site of hunger modulation to sensory neurons and two second-order neurons. Although caveats of artificial stimulation exist, the consistent changes seen across different light intensities for neural manipulation early in the pathway, but not downstream, argues that these results are robust. These studies demonstrate that hunger acts at a few critical nodes to modulate feeding initiation: sensory neurons increase detection sensitivity and second-order neurons amplify pathway activation. It will be interesting to examine whether hunger modulation of sensory and second-order neurons occurs independently or over different time scales to adjust behavioral responses as starvation increases. In addition, the specific hunger signals that act on central neurons and their mechanism of modulation may now be explored.

### Bitter compounds inhibit premotor neurons to prevent feeding initiation

While previous studies have demonstrated interactions between sweet and bitter taste modalities at the level of sensory neurons (Chu et al., 2014; French et al., 2015; Jeong et al., 2013; Meunier et al., 2003) and through feedback from the mammalian gustatory cortex (Jin et al., 2022), this study reveals a third circuit strategy for weighing sweet and bitter tastes: a local inhibitory network. Inhibitory interactions between bitter and sugar pathways at the level of premotor neurons provides an elegant strategy to weigh incoming sugar and bitter taste information and adjust behavioral probability. In addition, by blocking activity at specific muscles, bitter detection may specifically change behavior to direct the proboscis away from a hazardous food source. The existence of a local inhibitory circuit for bitter-sweet integration has been recently postulated based on studies of mammalian taste circuitry (Jin et al., 2022) and may be a shared strategy across species. These multiple circuit mechanisms for suppression of sweet attraction by bitter signals may reflect the evolutionary importance of robust bitter taste avoidance.

By examining a complete sensorimotor pathway, we elucidate how a complex nervous system orchestrates the decision to initiate feeding and illuminate central modules that integrate taste detection with internal state. These central controls afford independent amplification and suppression of feeding and stand apart from sensory modulation as mechanisms that dynamically tune behavior. As sensory modulation may suffer from finite amplification and incomplete suppression, central modulation provides a strategy to bypass those limits, allowing a broader range and different temporal dynamics of modulation.

## Acknowledgments

We thank Lori Horhor, Jolie Huang, Neil Ming, Vivian Nguyen, Andrea Sandoval, Neha Simha, Rivka Steinberg, and Parisa Vaziri for EM tracing contributions. Vivek Jayaraman provided unpublished fly lines used in this study. This work was supported by NIH R01DC013280 (K.S.), NIH F32DK117671 (G.R.S.) and NIH F32DC018225 (P.S.). Neuronal reconstruction for this project took place in a collaborative CATMAID environment in which 27 labs are participating to build connectomes for specific circuits. Development and administration of the FAFB tracing environment and analysis tools were funded in part by National Institutes of Health BRAIN Initiative grant 1RF1MH120679-01 to Davi Bock and Greg Jefferis, with software development effort and administrative support provided by Tom Kazimiers (Kazmos GmbH) and Eric Perlman (Yikes LLC). Peter Li, Viren Jain and colleagues at Google Research shared automatic segmentation (Li et al., 2019). We acknowledge the Princeton FlyWire team and members of the Murthy and Seung labs for development and maintenance of FlyWire (supported by BRAIN Initiative grant MH117815 to Murthy and Seung). We thank Drs. Stefanie Hampel, Jinseop Kim, Mala Murthy, Andrew Seeds, Sebastian Seung, Ibrahim Tastekin and Rachel Wilson and members of their laboratories for FlyWire tracing. We thank K. Eichler and members of the Connectomics Group in the Dept Zoology, University of Cambridge (G. Jefferis, M. Costa) for FlyWire tracing and sharing some neurons ahead of publication. Confocal imaging for triple intersection characterization was conducted at the CRL Molecular Imaging Center, supported by the Helen Wills Neuroscience Institute and NSF DBI-1041078. We would also like to thank Holly Aaron and Feather Ives for their microscopy training and assistance. Members of the Scott lab provided comments on the manuscript.

## Author Contributions

P.K.S. and G.R.S. conceived and performed experiments and analyzed the data under guidance from K.S. P.K.S. and S.E. performed EM analyses, P.K.S. and G.R.S. performed behavioral analyses, G.R.S. performed calcium imaging studies. G.R.S., B.J.D., and K.S. collaborated in the generation of split-Gal4 lines for these studies. P.K.S., G.R.S., and K.S. wrote the manuscript. B.J.D. and S.E. provided advice and guidance on the work.

## STAR METHODS

### RESOURCE AVAILABILITY

#### Lead contact

Further information and requests for resources and reagents should be directed to and will be fulfilled by the Lead Contact, Kristin Scott.

#### Materials availability

This study did not generate new unique reagents. All fly lines will be available upon request.

#### Data and code availability

Data reported in this paper will be shared by the lead contact upon request.

### EXPERIMENTAL MODEL AND SUBJECT DETAILS

#### Rearing conditions and strains

All experiments were performed in the fruit fly *Drosophila melanogaster*. The key resources table lists the transgenic lines used in this study. Flies were reared on standard cornmeal-yeast-molasses media at 25°C with 65% humidity and a 12hr: 12hr light: dark cycle unless stated otherwise. Flies for optogenetic experiments were raised on standard food in darkness. Upon eclosion, adult flies were collected and maintained on standard food supplemented with 0.4 mM all-trans-retinal in darkness prior to experiments. Adult mated female flies were used for all experiments.

### METHOD DETAILS

#### Electron Microscopy Neural Reconstructions

Second-order neurons were reconstructed in a serial section transmission electron volume (Full Adult Female Brain, Zheng et al., 2018) using the CATMAID software (Saalfeld et al., 2009). Fully manual reconstructions were generated by following the branches of the neuron and marking the center of each branch, thereby creating a “skeleton” of each neuron. In addition to fully manual reconstructions, segments of an automated segmentation (Li et al., 2020) were proofread and expanded to generate complete reconstructions. To specifically reconstruct second-order sugar neurons, two different methods were used. First, random presynapses of skeleton 7349219 (Engert et al., 2021) were chosen using the reconstruction sampler function of CATMAID and downstream partners were reconstructed. Second, large automatically generated fragments downstream of sugar GRN axons were found, and expanded. Chemical synapses were annotated as previously described (Zheng et al., 2018); specifically, at least three of four elements of a synapse were needed to call a synapse: a T-bar, postsynaptic density, synaptic vesicles, and a synaptic cleft. All reconstructions for which there is a corresponding split-Gal4 were assembled and proofread to near completion.

#### Flywire connectivity analysis

Neurons corresponding to those traced in CATMAID were located in Flywire (Flywire.ai); both reconstructions use the same underlying EM data (Zheng et al., 2018). To identify neurons upstream or downstream of a set of Flywire neurons, we used CAVE (connectome annotation versioning engine; Buhmann et al., 2021; Heinrich et al., 2018). To identify synapses of fairly high confidence, we chose a “cleft_score” cutoff of 100 (Heinrich et al., 2018).

The CATMAID skeleton IDs and Flywire IDs for each reconstructed neurons are listed here: Billiards (CATMAID: 8606542, Flywire: 720575940634231886), Bract1 (17024882, 720575940625204508), Bract2 (17542353, 720575940637873717), Clavicle (10150139, 720575940620111024), Dandelion (17249809, 720575940628601052), Fdg (16783943, 720575940632291554), FMIn (8952676, 720575940645551748), Fuchs (7929209, 720575940623691196), Fudog (7983275, 720575940630459463), G2N-1 (15079937, 720575940606258268), MN9 (16866694, 720575940616055252), Phantom (16762541, 720575940618879604), Quasimodo (8275570, 720575940619419814), Rattle (16238926, 720575940608777796), Rounddown (16886973, 720575940609112018), Roundup (16002203, 720575940620364549), Scapula (16887116, 720575940624539966), Specter (17579359, 720575940616547141), Sternum (17533840, 720575940643288356), Usnea (14890522, 720575940615947993), Zorro L (7574284, 720575940643219566), Zorro R (7899212, 720575940629888530). FAFB neuronal reconstructions will be available from Virtual Fly Brain (https://fafb.catmaid.virtualflybrain.org/).

#### Genetic access to Cleaver

To gain specific genetic access to Cleaver, we used a triple intersection approach. In this approach, CsChrimson-mVenus will only be expressed where the expression patterns of the AD, DBD, and LexA overlap. SS31022 (Sterne et al. 2021) labels both Cleaver and Usnea. To specifically access Cleaver, virgins of 20xUAS>dsFRT>csChrimson-mVenus;8XLexAop2-FLPL(attP40);Dfd-LexA were crossed to males of SS31022. To specifically access Usnea in SS31022, virgins of 20xUAS>dsFRT>csChrimson-mVenus;8XLexAop2-FLPL(attP40);Scr-LexA were crossed to males of SS31022. For each intersection, female progeny without balancers were selected for behavioral analysis.

To visualize triple intersection expression patterns, brains were dissected as described (https://www.janelia.org/project-team/flylight/protocols, ‘Dissection and Fixation 1.2% PFA’).

The following primary antibodies were used:

-1:40 mouse α-Brp (nc82) (DSHB, University of Iowa, USA)

-1:1000 chicken α-GFP (Invitrogen A10262) The following secondary antibodies were used:

-1:500 α-mouse AF647 (Invitrogen, A21236)

-1:1000 α-chicken AF488 (Life Technologies, A11039)

Immunohistochemistry was carried out as described (https://www.janelia.org/project-team/flylight/protocols, ‘IHC-Anti-GFP’) substituting the above antibodies and eschewing the pre-embedding fixation steps. Ethanol dehydration and DPX mounting was carried out as described (https://www.janelia.org/project-team/flylight/protocols, ‘DPX Mounting’). Images were acquired with a Zeiss LSM 880 NLO AxioExaminer. A Plan-Apochromat 25×/0.8 objective was used at zoom 0.7. Acquired images had a voxel size of 0.59 μm × 0.59 μm × 1.50 μm.

#### NBLAST Analysis

NBLAST analysis was used to match neurons reconstructed in EM to neurons labeled by split-GAL4 lines (Costa et al., 2014). Reconstructed neurons from CATMAID were transformed into the JRC2018U template space using NAVIS (Bates et al., 2020a; Schlegel, et al., 2021) and compared to a light-level library of 122 SEZ cell types in the SEZ split-GAL4 collection (Sterne et al., 2021). In addition, we added a representative image from a split-GAL4 we designed to cover a cell type reported here, Zorro, using previously described methods (Sterne et al., 2021). Each reconstructed neuron on the right of the brain was compared to every SEZ cell type in the library using the natverse toolkit in R (Bates et al., 2020). Normalized, mean scores were calculated to control for neuron size and segment number. The highest scoring light-level cell type for each reconstructed neuron was considered a match if the normalized, mean NBLAST score was greater than 0.4.

Reconstructed cell types with matches include the following *FAFB IDs* (Top match cell type, NBLAST score): *Bract1* (Bract, 0.58), *Bract2* (Bract, 0.57), *Clavicle* (Clavicle, 0.54), *Cleaver* (Cleaver, 0.57), *Fdg* (Fdg, 0.64), *FMIn* (FMIn, 0.61), *Fudog R* (Fudog, 0.43), *G2N-1* (G2N-1, 0.49), *Phantom* (Phantom, 0.67), *Rattle* (Rattle, 0.60), *Roundup* (0.63), *Usnea* (Usnea, 0.50), *Zorro R* (Zorro, 0.54). Reconstructed cell types which did not return matches include the following *FAFB IDs* (Top match cell type, NBLAST score): *Billiards* (Phantom, 0.17), *Buster* (Marge, 0.30), *Dandelion* (Clavicle, -0.16), *Fuchs* (Phantom, 0.37), *Quasimodo* (Puddle, 0.32), *Rounddown* (Roundup, 0.28), *Roundtree* (Puddle, 0.27), *Specter* (Usnea, 0.16), *Sternum* (Rattle, 0.10).

#### Optogenetic Activation

PER was scored as previously described (Mann, Gordon and Scott, 2013). Female flies were raised on standard cornmeal-yeast-molasses medium, until 48 hours before experiments, when flies were placed on molasses food with 0.4 mM retinal. Three to five-day-old flies were anesthetized with carbon dioxide, mounted onto a glass slide with nail polish, and allowed to recover for two hours in a humidified chamber at 22°C. For experiments in Figure 2A, flies were activated with 153 uW/mm^2^ 635 nm laser light (Laserglow). Flies were scored for whether they extended their proboscis within a 5 second period in response to light.

For food-deprivation experiments, flies were raised as above, except 48 hours before experiments, flies were wet-starved by placing them in a vial with a water saturated kimwipe supplemented with .4 mM retinal. Flies were activated with a 635 nm laser at four different light intensities: 1.8, 8.9, 17.8 and 153 uW/mm^2^.

#### GtACR1 silencing

Three-day-old female flies were raised on standard food, and transferred to standard food with 0.4 mM all-trans retinal for two days. Next, flies were wet-starved with 0.4 mM retinal in water for 24 hours. Flies were anesthetized with carbon dioxide, mounted onto a glass slide with nail polish, and allowed to recover for two hours in a humidified chamber at 22°C. A green laser (532 nm, LaserGlow LBS-532) was used to acutely silence neurons using GtACR1 (Mohammad et al., 2017). Flies were water satiated, then presented with either 50 mM sucrose or 100 mM sucrose three times to the proboscis, and the number of flies that extended at least once were recorded.

#### *In vivo* sample preparation for calcium Imaging

Mated female flies were dissected for calcium imaging studies 14 to 21 days post-eclosion as previously described (Harris et al., 2015) with the following modifications. Flies were briefly anesthetized with ice as they were placed in a custom plastic holder at the cervix to isolate the head from the rest of the body. Then, the head was then immobilized using UV glue. To provide unobstructed imaging access to the SEZ, the esophagus was cut. Flies in fed, food-deprived, desiccated, and thristy-like (pseudodessicated) conditions were generated as follows:

Fed: Flies were placed in a fresh vial containing standard cornmeal-yeast-molasses media 18-24 hours prior to imaging. Following dissection, samples were bathed in ∼250 mOsmo Artificial Hemolymph-Like solution (AHL) (“artificial hemolymph”) and imaged immediately.

Food-deprived: Flies were food deprived in a vial containing a wet kimwipe for 18-24 hours prior to imaging. Following dissection, samples were bathed in ∼250 mOsmo AHL and imaged immediately.

Desiccated: Flies were placed in a vial containing 5 grams of Drierite™ for two hours. A cotton ball was used to isolate flies from the desiccant inside the vial, and the vial was closed with parafilm to create a dry chamber. Following dissection, samples were bathed in ∼250 mOsmo AHL and imaged immediately. Hemolymph signals of thirst, such as osmolality, may be perturbed in our calcium imaging studies, limiting our ability to accurately assess a thirsty state (Jourjine et al., 2016).

Thirsty-like (Pseudodessicated): Flies were placed in a fresh vial containing standard cornmeal-yeast-molasses media 18-24 hours prior to imaging. Following dissection, samples were bathed in ∼350 mOsmo AHL (“high osmolality artificial hemolymph”) and allowed to rest for one hour prior to imaging.

#### Calcium imaging with taste stimulation

For imaging responses to taste solutions, females of UAS-CD8- tdTomato;20XUAS-IVS-GCaMP6s(attP5);20XUAS-IVS-GCaMP6s(VK00005) were crossed to males for each split-GAL4 line, and female progeny without balancers were selected for imaging. The following tastants were used: double-distilled water (“water”), 1M sucrose (“sugar”), or 10 mM denatonium plus 100 mM caffeine in 20% polyethylene glycol (PEG) (“bitter”). Taste solutions were delivered to the proboscis using a glass capillary (1.0 mm OD/ 0.78 mm ID) filled with ∼4 µL of taste solution and positioned at the tip of the proboscis using a micromanipulator. Taste solutions were drawn away from the tip of the capillary at the beginning of each imaging trial using slight suction generated by an attached 1 mL syringe, and delivered to the proboscis at the relevant time during imaging with light pressure applied to the syringe.

Calcium imaging was performed using either a 1-photon or 2-photon microscope.

For cell types in close proximity to the surface of the SEZ, 1-photon imaging was performed using a 3i spinning disc confocal microscope with a piezo drive and a 20x water immersion objective (NA=1.0) with a 2.5x magnification changer. 55 frames of 8 z sections spaced at 1 micron intervals were binned 4x4 and acquired at 0.8 Hz using a 488 nm laser. Taste solutions were in contact with the proboscis labellum from frame 20 to frame 25. Cell types imaged using a 1-photon microscope are Clavicle, Fdg, FMIn, G2N-1, Phantom, Usnea, and Zorro. For cell types that arborize deeper in the SEZ, 2- photon imaging was performed using a Scientifica Hyperscope with resonant scanning, a piezo drive, and a 20x water immersion objective (NA=1.0) with 4x digital zoom. 80 stacks of 20 z sections spaced at 2 micron intervals were acquired at 0.667 Hz using a 920 nm laser. Taste solutions were in contact with the proboscis labellum from frame 30 to frame 40. Cell types imaged using a 2-photon microscope are Bract, Rattle, and Roundup.

#### Calcium imaging with optogenetic activation of gustatory receptor neurons

For imaging responses in the Fdg cell type to optogenetic activation of gustatory receptor neurons, females of 13XLexAop2-IVS-p10-ChrimsonR-mCherry(attP18);Gr5a- LexA;20XUAS-IVS-jGCaMP7b(VK00005), 13XLexAop2-IVS-p10-ChrimsonR-mCherry(attP18);20XUAS-IVS-jGCaMP7b(attP5); ppk28-LexA, or 13XLexAop2-IVS- p10-ChrimsonR-mCherry(attP18);20XUAS-IVS-jGCaMP7b(attP5); Gr66a-LexA were crossed to males of SS46913 (Sterne et al. 2021). For sugar and bitter integration experiments, virgins of a stock composed of either SS47082 (G2N-1) or SS47744 (Roundup) and 20xUAS-IVS-Syn21-Syt::Op-jGCaMP7b(attP18) were crossed to males of 13XLexAop2-IVS-p10-ChrimsonR-mCherry(attP18); Gr5a-LexA::VP16(12-1);, 13XLexAop2-IVS-p10-ChrimsonR-mCherry(attP18);Gr5a-LexA::VP16(12-1); Gr66a- LexA, or 13XLexAop2-IVS-p10-ChrimsonR-mCherry(attP18);;Gr66a-LexA; and female progeny without balancers were selected for imaging. 2-photon imaging was performed as described above for imaging with taste stimulation, but 660 nm light was used to activate gustatory receptor neurons in place of direct stimulation of the proboscis with taste solutions. Two-second light pulses were delivered three times at 10-second intervals during imaging, and light was delivered through the objective in a widefield fashion under the control of a custom ScanImage plugin.

#### Calcium imaging analysis

Image analysis was carried out in Fiji (Schindelin et al., 2012), CircuitCatcher (Bushey 2021), Python, and R. First, in Fiji, Z stacks for each time point were maximum intensity projected and then movement corrected using the StackReg plugin with ‘Rigid Body’ or ‘Translation’ transformation (Thevenaz et al. 1998). Next, using CircuitCatcher, an ROI containing the neurites of the cell type of interest was selected along with a background ROI, and average fluorescence intensity for each ROI at each timepoint was retrieved. Then, in Python, background subtraction was carried out for each timepoint (Ft). To calculate Finitial, initial fluorescence intensity was calculated as the mean corrected average fluorescence intensity from frame 9 to 18 (for 1-photon imaging) or frame 0 to 19 (for 2-photon imaging and optogenetic imaging). Finally, the following formula was used to calculate ΔF/F: Ft-Finitial/Finitial. Area under the curve was approximated with the trapezoidal rule in Python using the NumPy.trapz function. Area under the curve was assessed from frames 20 to 25 (for 1-photon imaging), from frames 30 to 40 (for 2- photon imaging with taste stimulation), from frames 15 to 18 (for 2-photon imaging with optogenetic activation).

### QUANTIFICATION AND STATISTICAL ANALYSIS

Statistical tests for behavioral assays were performed in Prism. For analysis of Proboscis Extension Response Assays, Fisher’s Exact Test was used in comparing the fraction of PER responses in experimental versus control flies. Statistical analysis of calcium imaging was carried out in R and Python. For imaging experiments carried out in a block design with three treatments (Figure 4, Figure S4), Quade tests were carried out in R using the PMCMRplus package (Thorsten 2021). Quade test was chosen because it is more powerful than Friedman for a block-design experiment with three treatments (Conovoer, 1999). Other statistical analyses of calcium imaging were carried out in Python using the SciPy (Virtanen et al. 2020) and scikit-posthocs packages (Therpilowski 2019).

---

**Figure S1.**
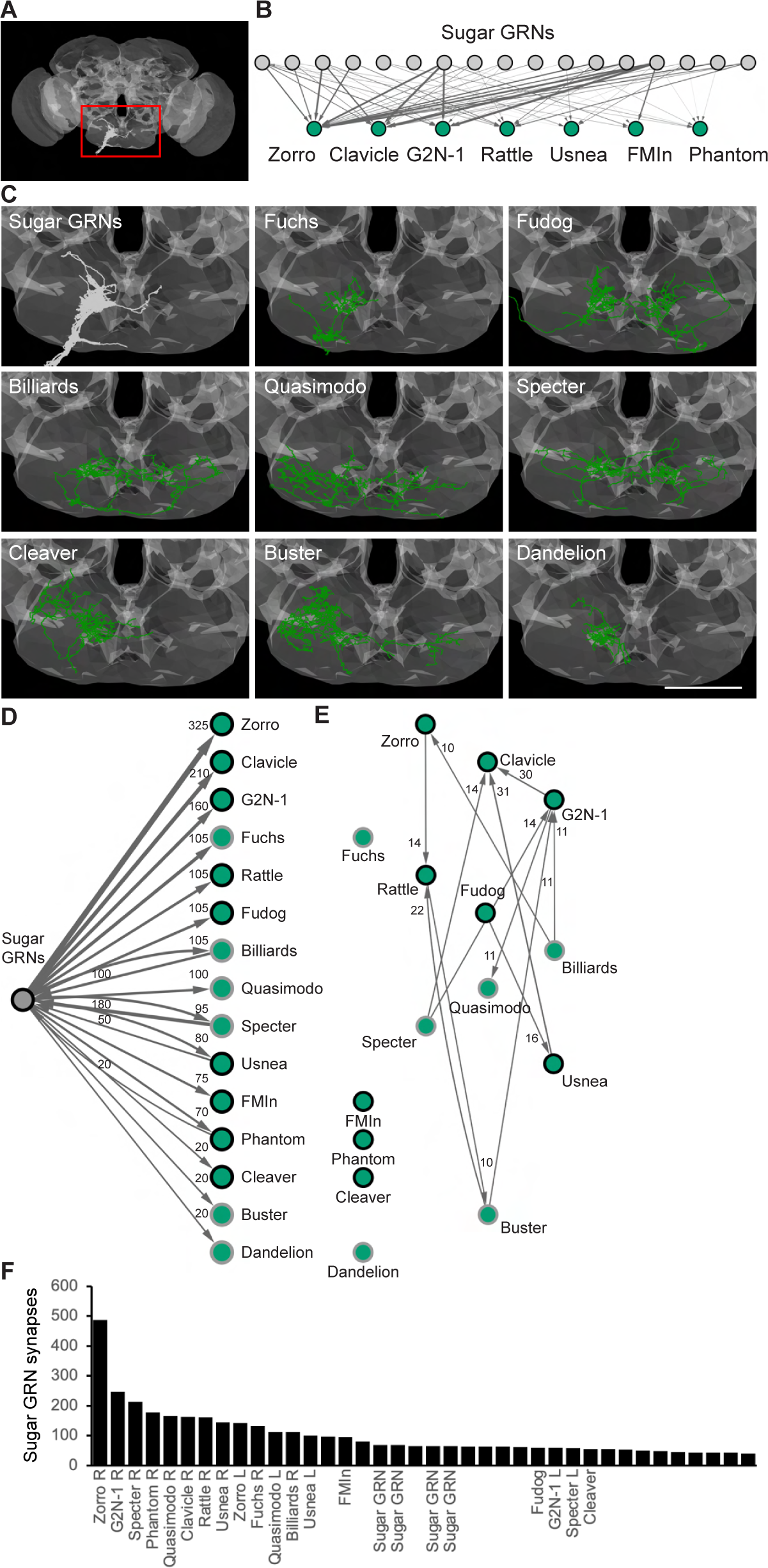
Anatomy of reconstructed second-order neurons and their connectivity, Related to Figure 1. (A) Schematic of the entire fly brain, showing EM reconstructed sugar-sensing gustatory receptor neurons (GRNs) in gray. FAFB neuropil space is shown in darker gray. Area outlined in red is enlarged in the first panel of C. (B) Synaptic connectivity of 17 previously identified candidate sugar GRNs onto second-order neurons that elicit PER. Line thickness represents the number of synapses, with a minimum of 6 synapses to a maximum of 46 synapses. (C) Anatomy of second-order candidate sugar EM reconstructed neurons. Scale bar is 50 μm. (D) Synaptic connectivity from sugar GRNs onto and from second-order neurons. Second-order neurons identified by EM and present in a split-Gal4 line (black circles); second-order neurons identified by EM only (gray circles). (E) Synaptic connectivity between second-order neurons. (F) Neurons with the most synapses from 17 candidate sugar GRNs based on Flywire predicted synapses (n ≥40), with x-axis labelling neurons identified in this study.

**Figure S2.**
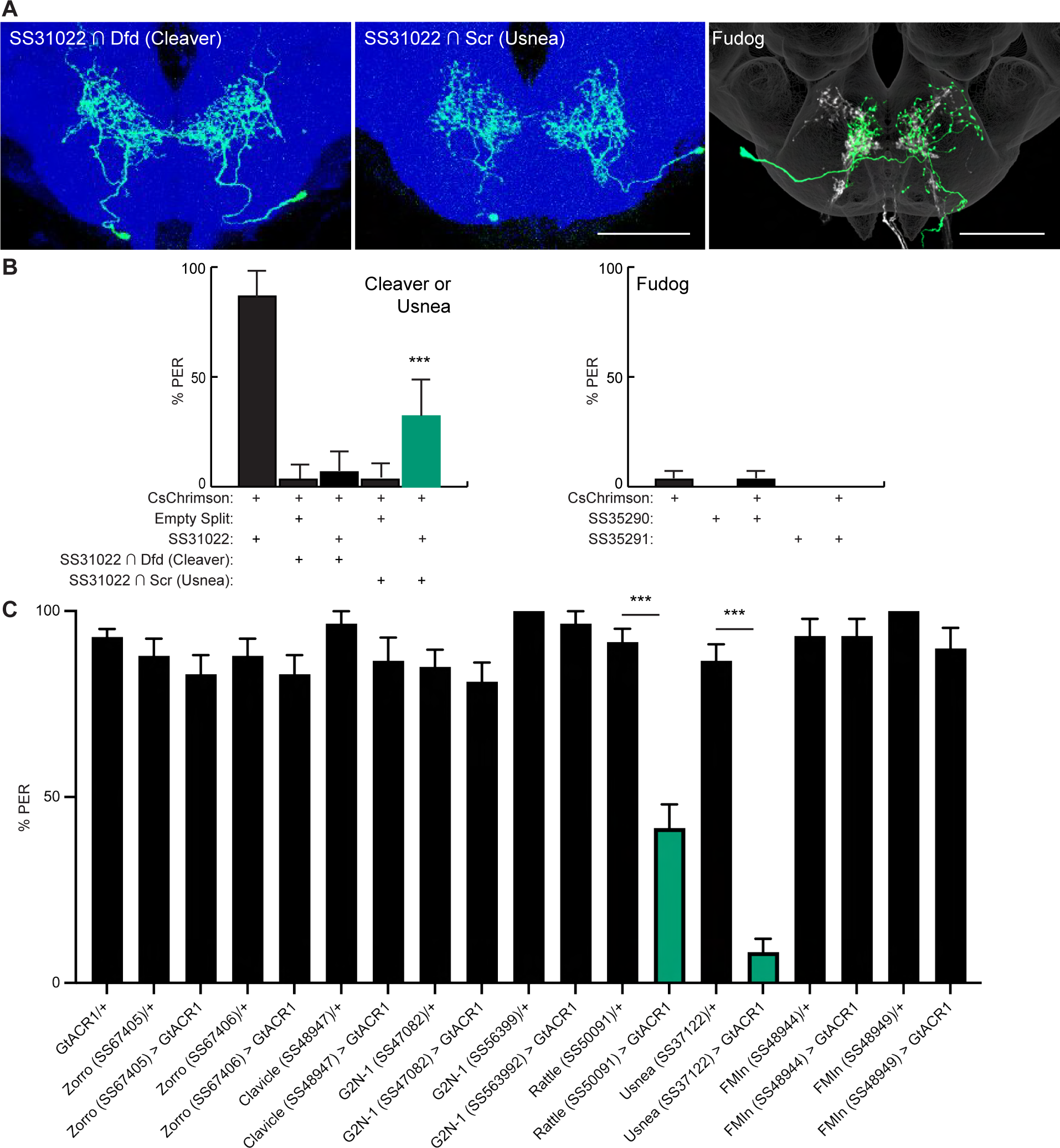
Additional proboscis extension phenotypes of second order neurons, Related to Figure 2. (A) Light microscopy images of Cleaver, Usnea and Fudog. Specific lines for Cleaver and Usnea were generated using a triple-intersection approach (see Methods). In the Fudog image, sugar GRNs are depicted in white. Scale bar is 50 μm. JRC 2018 unisex coordinate space shown in blue (Cleaver and Usnea) or dark gray (Fudog). (B) Activation of Usnea, but not Cleaver or Fudog, elicits proboscis extension. (C) Hyperpolarization of Rattle and Usnea inhibited proboscis extension to 100 mM sucrose, but hyperpolarization of other second-order neurons did not.

**Figure S3.**
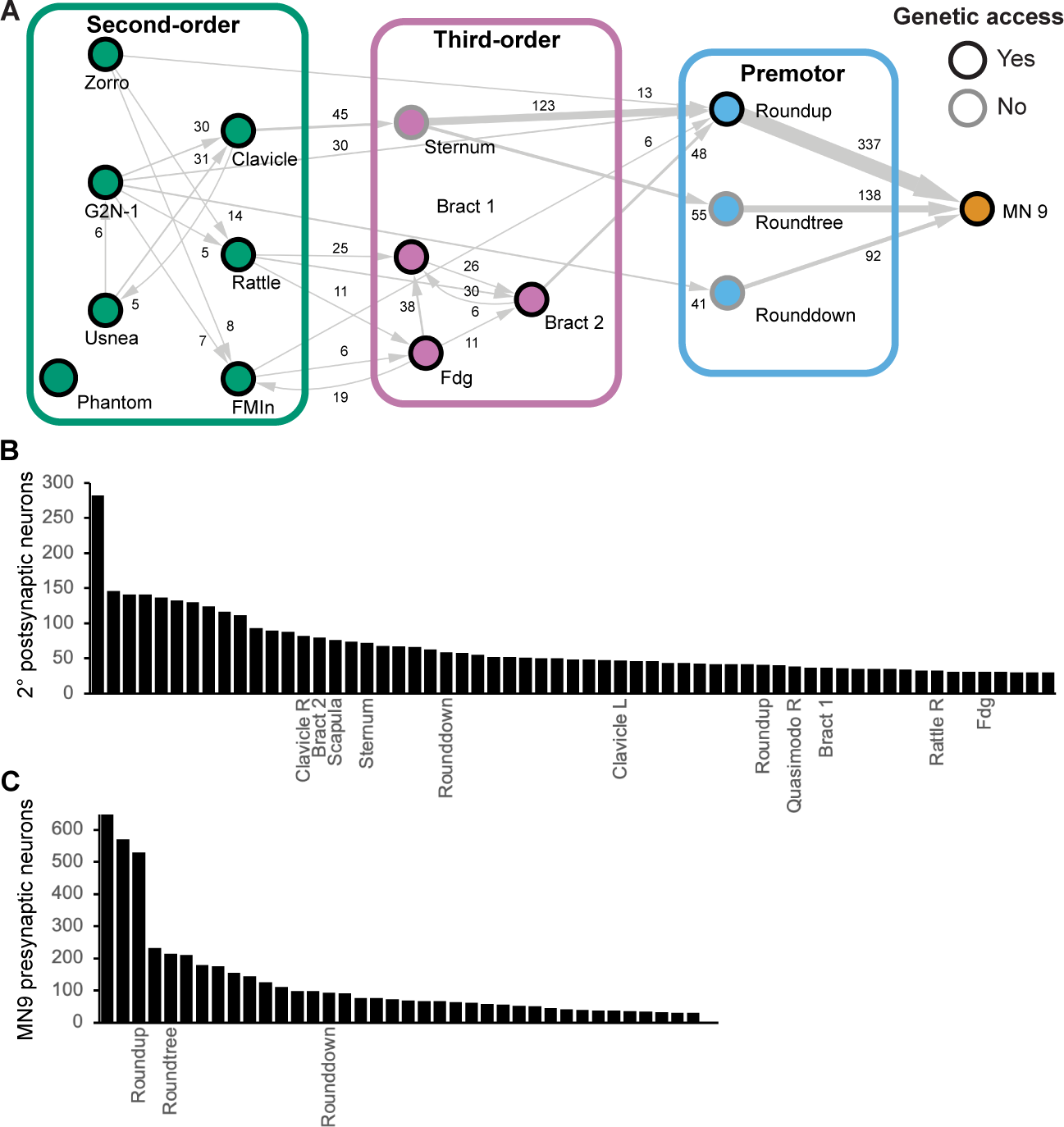
Synaptic connectivity in the feeding initiation circuit, Related to Figure 3. (A) Schematic of the feeding initiation circuit, with circles outlined in black for neurons with split- Gal4 lines, circles outlined in gray for neurons without split-Gal4 lines. Line thickness represents connectivity of more than 5 synapses, synapse numbers labelled. (B) Neurons with the most synapses from second order neurons that elicit PER. based on Flywire predicted synapses **(n**≥30) x-axis labels neurons identified in this study. (C) Neurons with the most synapses onto MN9, ased on Flywire predicted synapses (n≥30) with x-axis labels neurons identified in this study.

**Figure S4.**
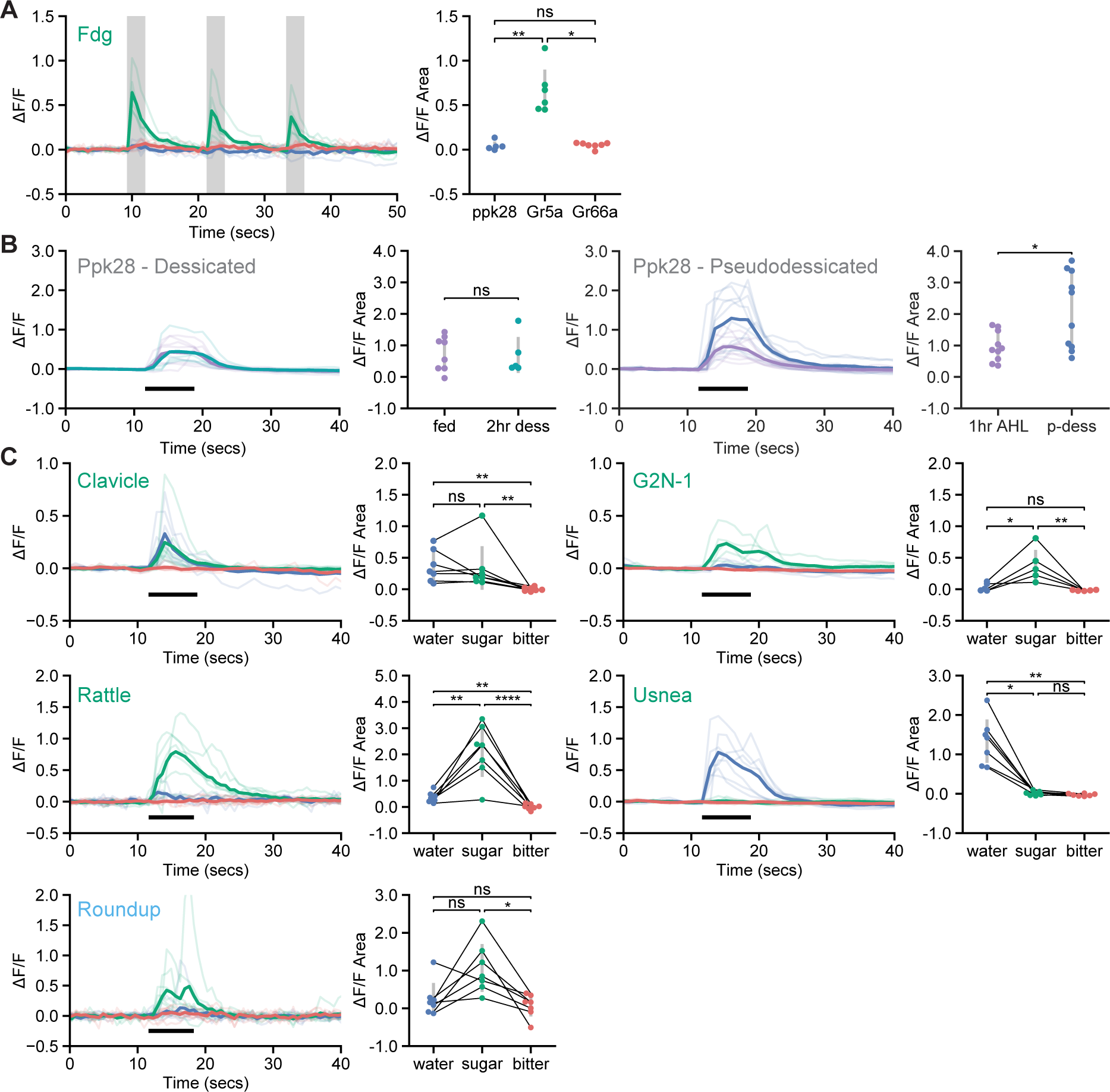
Taste responses of feeding initiation neurons, Related to Figure 4. (A) Calcium responses of Fdg to optogenetic activation of water (blue, ppk28-LexA), sugar (green, Gr5a-LexA), or bitter (red, Gr66a-LexA) GRNs in food-deprived flies. To examine Fdg responses, GCaMP7b was expressed using SS46913. Fluorescence traces are shown on the left of the panel (ΔF/F), while ΔF/F area for each trace is shown on the right. Stimulation with 660 nm light is indicated with vertical gray bars. Kruskal Wallace test with Dunn’s test using Holm’s correction to adjust for multiple comparisons, ns p>0.05, *p≤0.05, **p≤0.01. (B-C) Calcium responses of water gustatory sensory neurons and feeding initiation neurons to taste stimulation of the proboscis. GCaMP6s fluorescence traces are shown on the left of each panel (ΔF/F), while ΔF/F area for each trace is shown on the right. Significance levels: ns p>0.05, *p≤0.05, **p≤0.01, ***p≤0.001, ***p≤0.0001. (B) Taste responses to water in fed (purple) versus two hour desiccated (2hr dess, teal) flies (left). Taste responses to water in fed flies allowed to rest for one hour in low osmolality artificial hemolymph after dissection before imaging (1hr fed, purple) versus pseudodessicated, thirsty-like flies (p-dess, blue) (right). (C) Taste responses in pseudodessicated, thirsty-like flies. The proboscis of each tested individual was stimulated with water (green), sugar (blue), and bitter (red) tastants in sequential trials during the indicated period (thick black line). Thin black lines indicate sample pairing. The following split-GAL4 lines were imaged for each cell type: Clavicle; SS48947, G2N-1; SS47082, Usnea; SS37122, Rattle; SS50091, Roundup; SS47744. Quade’s Test with Quade’s All Pairs Test, using Holm’s correction to adjust for multiple comparisons.

**Figure S5.**
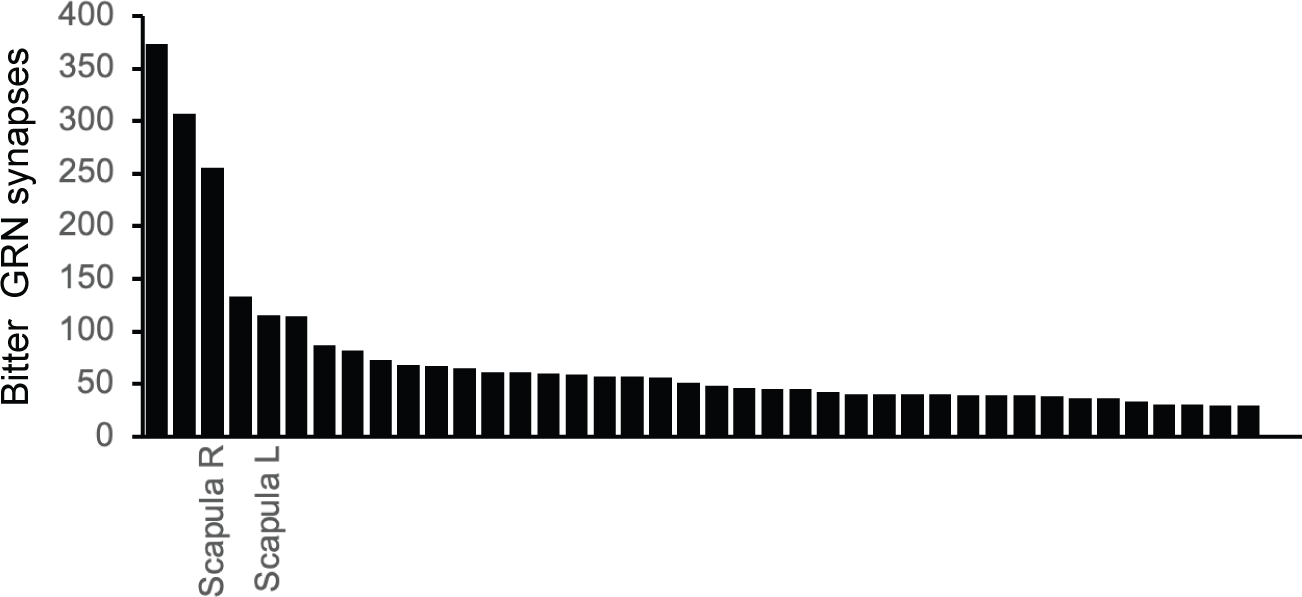
Second-order bitter neurons, related to Figure 6. Neurons with the most synapses from candidate bitter GRNs based on Flywire predicted synapses (n≥30) with x-axis labelling neurons identified in this study.

